# Intravenous haloperidol and cocaine alter the distribution of T CD4^+^ and B lymphocytes and NKT cells in rats

**DOI:** 10.1101/2021.11.28.470229

**Authors:** Maciej M. Jankowski, Bogna M. Ignatowska-Jankowska, Wojciech Glac, Marek Wiergowski, Paulina Kaźmierska-Grębowska, Artur H. Świergiel

## Abstract

Modulation of dopamine transmission evokes strong behavioral effects that can be achieved by psychoactive drugs such as haloperidol or cocaine. Cocaine non-specifically increases dopamine transmission by blocking dopamine active transporter (DAT) and evokes behavioral arousal, while haloperidol is a non-specific dopamine D2 receptor antagonist with sedative effects. Interestingly, dopamine has been found to affect immune cells in addition to its action in the central nervous system. Here we address the possible interactions between haloperidol and cocaine and their effects on both immune cells and behavior in freely moving rats. We use an intravenous model of haloperidol and binge cocaine administration to evaluate the drugs’ impact on the distribution of lymphocyte subsets in both the peripheral blood and the spleen. We assess the drugs’ behavioral effects by measuring locomotor activity. Cocaine evoked a pronounced locomotor response and stereotypic behaviors, both of which were completely blocked after pretreatment with haloperidol. The results suggest that blood lymphopenia which was induced by haloperidol and cocaine (except for NKT cells), is independent of dopaminergic activity and most likely results from the massive secretion of corticosterone. Haloperidol pretreatment prevented the cocaine-induced decrease in NKT cell numbers. On the other hand, the increased systemic dopaminergic activity after cocaine administration is a significant factor in retaining T CD4^+^ and B lymphocytes in the spleen.

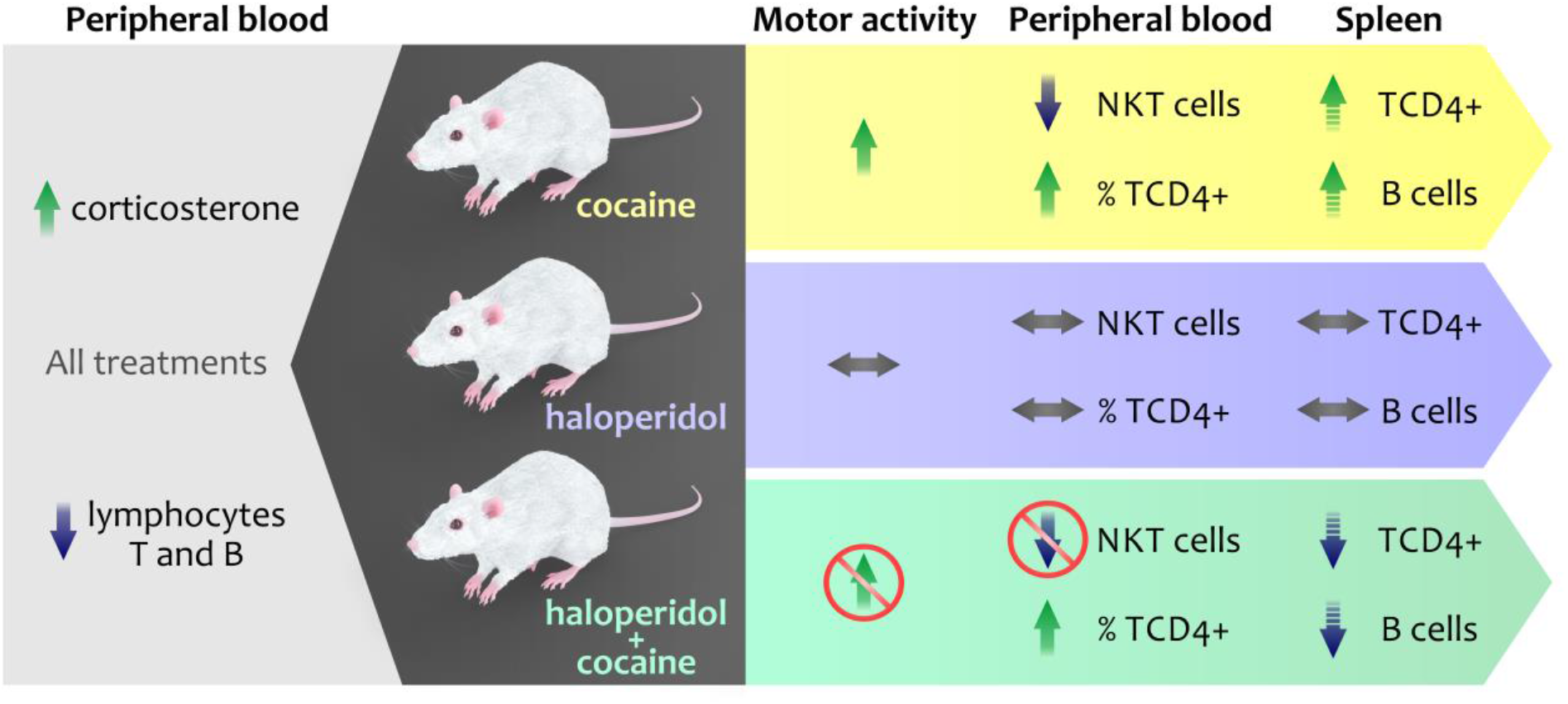

## 1. Introduction

Haloperidol is a potent neuroleptic drug frequently used to treat psychotic episodes in humans [1,2]. It is a non-specific D2 dopamine (DA) receptor antagonist that induces strong behavioral effects, including reduced psychomotor activity and catalepsy [3–5]. Haloperidol can affect neurocognitive performance [6,7] and the immune system [1,8–11]. In the brain itself, however, the effect of haloperidol on local immunocompetent cells is still a matter of debate [12–14].

Cocaine suppresses the immune system via both central and peripheral mechanisms [15–18]. The hypothalamic-pituitary-adrenal (HPA) axis and sympathetic nervous system are the primary pathways through which cocaine modulates the immune system [17,19,20]. However, in addition to neuroendocrine and sympathetic activation, cocaine may affect lymphocytes by increasing DA signaling at both central and peripheral sites. At the central site, cocaine can affect the immune system via alterations in DA transmission in specific brain regions [21].

At the periphery, both cocaine and haloperidol can directly modulate DA signaling between lymphocytes in peripheral blood, or in lymphatic organs and thereby change their activity, selective adhesion, migration, and homing [22,23]. DA receptor gene polymorphisms have been found to alter the count of circulating T lymphocytes [24]. Increased D3 DA receptor mRNA levels in circulating T lymphocytes were proposed as a potential diagnostic biomarker of psychotic disorders [25]. Human and rat lymphocytes express tyrosine hydroxylase as well as D1- and D2-like DA receptors [25–28]. The dopamine active transporter (DAT) and vesicular monoamine transporters (VMAT) have been found in human and rat lymphocytes [29,30]. Rat lymphoid tissues, such as the spleen and thymus, express DA, D1- and D2-like DA receptors, DAT, and VMAT [30–33]. Considering this accumulating evidence, we hypothesized that DA signaling affected by haloperidol and cocaine may partially mediate the effects of those drugs on the distribution of lymphocyte subsets in the peripheral blood and spleen.

We tested our hypothesis in freely moving rats using intravenous haloperidol and binge cocaine administration [34]. We measured locomotor activity during and after cocaine infusions to assess the behavioral effects of the drugs. Under these conditions, we determined the effects of haloperidol and cocaine on the numbers and proportions of T CD3^+^, T CD4^+^, T CD8^+^, B CD45RA^+^, Natural Killer (NK) CD161a^+^, and NKT CD3^+^CD161a^+^ cells in the peripheral blood and spleen. Additionally, we measured plasma corticosterone concentration to evaluate the HPA axis activation. Serum concentrations of IL-4 and IFN-γ were chosen as *in vivo* markers of anti- and pro-inflammatory cytokine profiles, respectively. Finally, we measured serum concentrations of cocaine and its metabolites (namely, benzoylecgonine and ecgonine methyl ester) to determine whether the binge cocaine administration resulted in the accumulation of cocaine or of its metabolites in the blood.

## 2. Results

### 2.1. Locomotor activity

Cocaine infusions significantly increased locomotor activity (including stereotyped movements) during the 90-minute session (F (1,28) = 75.1, p < 0.0001). This effect was completely blocked by pretreatment with haloperidol (Figure 1A-C, cocaine × haloperidol interaction: F (1,28) = 72.4, p < 0.0001; t (1,14) = 8.72, p < 0.0001). The separate measurements for each 30-minute interval between cocaine or saline infusions revealed that haloperidol alone significantly reduced locomotor activity (compared to vehicle-treated rats), but only during the first 30 minutes (haloperidol: t (1,14) = 3.2, p < 0.05, data not shown).

**Figure 1.**
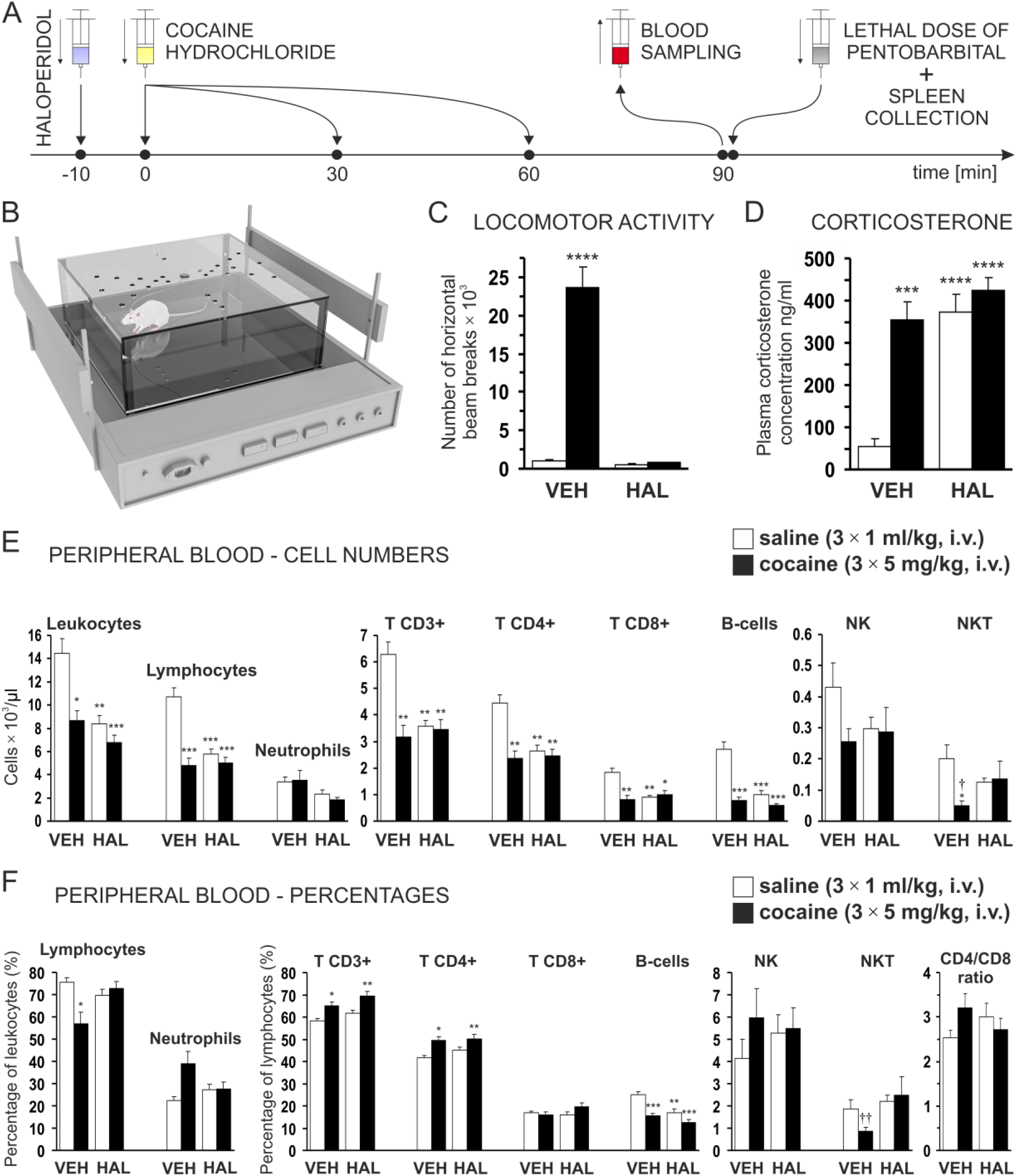
(A) Drug administration schedule with blood and spleen sampling times. (B) Experimental setup for multiple intravenous infusions and blood sampling in freely moving rats. (C-F) Effects of haloperidol (HAL) and cocaine i.v. infusions on (C) locomotor activity, (D) plasma corticosterone concentration, (E-F) leukocyte, lymphocyte, neutrophil, and lymphocyte subsets (E) cell numbers and (F) percentages and CD4/CD8 ratio in peripheral blood. HAL (1 mg/kg) or vehicle (VEH; 1 ml/kg) were administered 10 minutes before the first infusion of either cocaine (3 × 5 mg/kg, at 30-minute intervals) or saline (3 × 1 ml/kg, at 30-minute intervals). Each group represents the mean ± SEM of 8 animals. Significant differences compared to the saline-vehicle treated control rats are indicated as follows: *p<0.05, **p<0.01, ***p<0.001, ****p<0.0001 and compared to rats treated with haloperidol alone, where †p<0.05, ††p<0.01.

### 2.2. Plasma corticosterone

Administration of cocaine and haloperidol significantly increased plasma corticosterone concentrations (Figure 1D, cocaine: F (1,28) = 30.9, p < 0.0001, t (1,14) = 6.4, p < 0.001; haloperidol: F (1,28) = 25.2, p < 0.0001, t (1,14) = 7.1, p < 0.0001; cocaine × haloperidol interaction: F (1,28) = 12.8, p < 0.01, haloperidol + cocaine: t (1,14) = 10.7, p < 0.0001).

### 2.3. Lymphocytes in peripheral blood

Intravenous cocaine or haloperidol alone significantly reduced total leukocyte numbers in peripheral blood (cocaine: F (1,28) = 18.1, p < 0.001, t (1,14) = 3.8, p < 0.05; haloperidol: F (1,28) = 20.8, p < 0.0001, t (1,14) = 4.3, p < 0.01). Likewise, cocaine administered after haloperidol significantly reduced leukocyte numbers (interaction cocaine × haloperidol: F (1,28) = 5.5, p < 0.05, t (1,14) = 5.5, p < 0.001). The decrease in leukocyte numbers induced by cocaine alone resulted mainly from the reduction in lymphocyte numbers (F (1,28) = 33.2, p < 0.0001, t = 6.1, p < 0.001), while the neutrophil numbers were not affected (Figure 1E). Cocaine administered following pretreatment with haloperidol decreased lymphocyte numbers to a similar degree as for cocaine alone (interaction cocaine × haloperidol: F (1,28) = 18.7, p < 0.001), but not selectively because haloperidol also decreased neutrophil numbers (haloperidol: F (1,28) = 6.8, p < 0.05). Lymphocyte depletion after cocaine and/or haloperidol treatment was similar and involved mainly T CD3+, T CD4+, T CD8+, and B (CD45RA+) lymphocytes (Figure 1E, interactions cocaine × haloperidol for the listed lymphocyte subsets: F (1,28) = 14.4, p < 0.001; 12.6, p < 0.01; 14.0, p < 0.001; and 23.0, p < 0.0001, respectively). NKT (CD3+CD161a+) cell numbers were significantly reduced by cocaine alone (cocaine: t (1,14) = 3.2, p < 0.05), and that effect was prevented by pretreatment with haloperidol (interaction cocaine × haloperidol: F (1,28) = 4.8, p < 0.05). NK (CD161a+) cell numbers were not significantly decreased following administration of cocaine and haloperidol (Figure 1E).

Cocaine treatment alone decreased the percentage of circulating lymphocytes compared to vehicle-treated control rats, and this effect was blocked by haloperidol (interaction cocaine × haloperidol: F (1,28) = 9.4, p < 0.01). The percentage of neutrophils was not significantly increased after treatment with cocaine alone (Figure 1F). Cocaine alone and cocaine after pretreatment with haloperidol significantly increased the percentage of T CD3^+^ lymphocytes (cocaine: F (1,28) = 18.3, p < 0.001; haloperidol: F (1,28) = 5.4, p < 0.05), and the effects of cocaine were not modified by haloperidol (F (1,28) = 0.07, p = 0.79). An increase in the proportion of T CD3^+^ cells after cocaine administration was induced by the significant increase in the percentage of T CD4^+^ lymphocytes (cocaine: F (1,28) = 16.7, p < 0.001), while the proportion of T CD8+ lymphocytes remained unchanged significantly (Figure 1F). Haloperidol and/or cocaine significantly reduced the percentages of B lymphocytes compared to control animals (cocaine: F (1,28) = 27.6, p < 0.0001; haloperidol: F (1,28) = 16.4, p < 0.001), and the effects of cocaine were not modified by haloperidol (interaction cocaine × haloperidol: F (1,28) = 3.8, p = 0.06). The percentage of NKT cells was decreased by cocaine (t (1,14) = 4.1, p < 0.01), and the effect blocked by haloperidol (Figure 1F, interaction cocaine × haloperidol not significant due to considerable variation: F (1,28) = 1.7, p = 0.19). The CD4/CD8 ratio was not significantly increased after cocaine alone.

### 2.4. Lymphocytes in the spleen

Cocaine alone and cocaine infused after blocking D2-like DA receptors with haloperidol had opposite effects on the numbers of splenocytes: cocaine alone (non-significantly) increased the numbers of splenocytes compared to control rats while cocaine infused after haloperidol (non-significantly) decreased the numbers of splenocytes. Consequently, the numbers of splenocytes in rats receiving infusions of cocaine following the pretreatment with haloperidol were significantly lower than in rats treated with cocaine alone (Figure 2, haloperidol: F (1,28) = 4.8, p < 0.05; interaction cocaine × haloperidol: F (1,28) = 6.2, p < 0.05; t (1,14) = 3.1, p < 0.05). The splenocyte numbers were affected mainly by changes in the numbers of lymphocytes (haloperidol: F (1,28) = 6.0, p < 0.05, cocaine × haloperidol interaction: F (1,28) = 8.1, p < 0.01; t (1,14) = 3.5, p < 0.05), while neutrophil numbers remained unchanged. The lymphocyte numbers were affected mainly by alterations in T CD3^+^, T CD4^+^ and B lymphocyte numbers (cocaine × haloperidol interactions: F (1,28) = 4.3, p < 0.05; F (1,28) = 7.3, p < 0.05; F (1,28) = 7.0, p < 0.05, respectively). As a result, the numbers of T CD3^+^CD4^+^ and B CD45RA^+^ lymphocytes were significantly lower in rats receiving cocaine after pretreatment with haloperidol than in rats treated with cocaine alone (t (1,14) = 3.3, p < 0.05; t (1,14) = 3.6, p < 0.05, respectively). The T CD8^+^, NK, and NKT cell numbers were non-significantly changed by the treatments (Figure 2).

**Figure 2.**
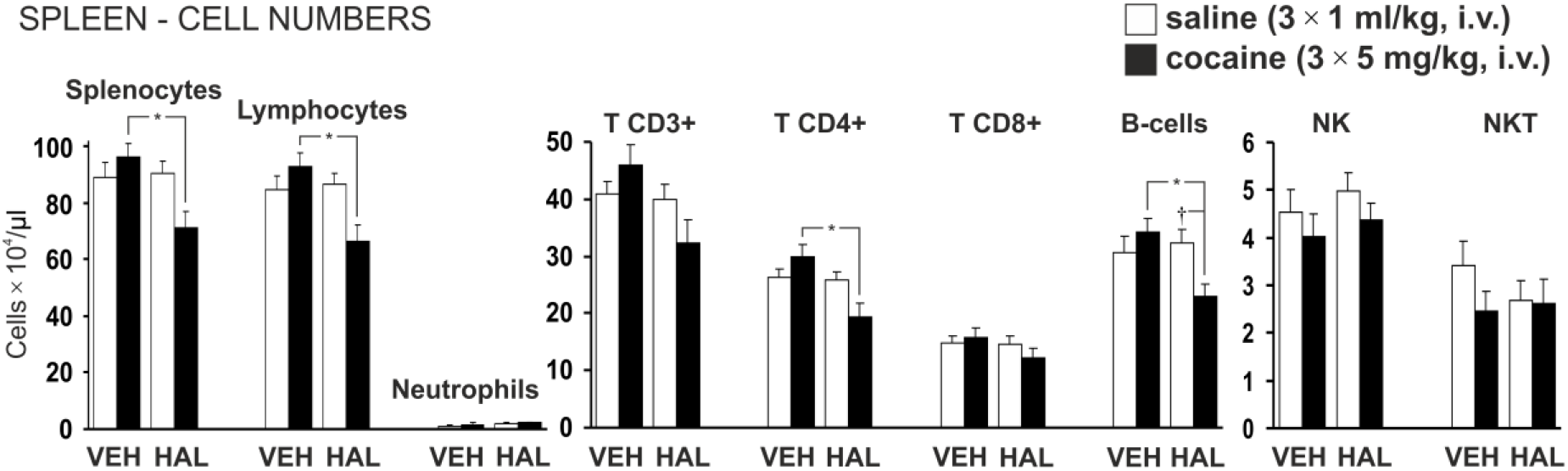
Effects of haloperidol (HAL) and cocaine intravenous administration on splenocyte, lymphocyte, neutrophil, and lymphocyte subsets numbers in the spleen of rats. Haloperidol (HAL; 1 mg/kg) or vehicle (VEH; 1 ml/kg) were administered 10 minutes before the first infusion of either cocaine (3 × 5 mg/kg, at 30-minute intervals) or saline (3 × 1 ml/kg, at 30-minute intervals). Each group represents the mean ± SEM of 8 animals and significant differences are indicated with, *p<0.05 and †p<0.053.

Proportions between T CD3^+^, T CD4^+^, T CD8^+^, and B lymphocytes in the rat spleen remained unchanged following all treatments (data not shown). The percentages of NK and NKT cells were decreased by cocaine. For NKT cells the effects were reversed by haloperidol (cocaine × haloperidol interaction: F (1,28) = 6.8, p < 0.05).

### 2.5. Serum IL-4 and IFN-γ concentrations

Serum concentrations of IL-4 and IFN-γ were non-significantly decreased after haloperidol and/or cocaine infusions (Figure 3).

**Figure 3.**
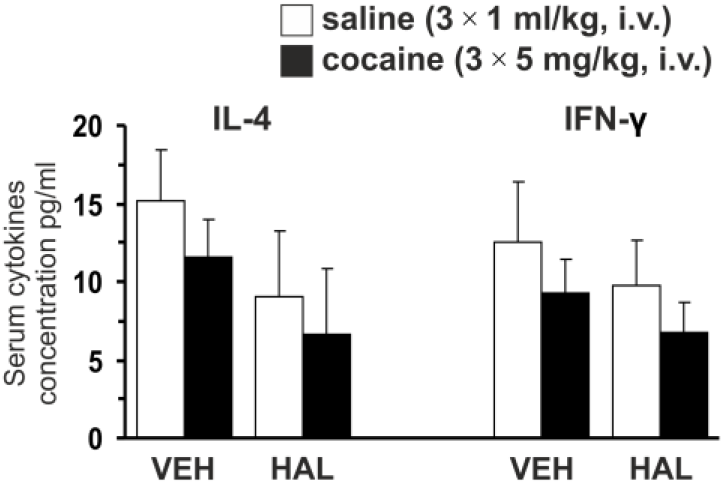
Serum concentrations of IL-4 and IFN-γ. Haloperidol (HAL; 1 mg/kg, i.v.) or vehicle (VEH; 1 ml/kg, i.v.) were administered 10 minutes before the first infusion of either cocaine (3 × 5 mg/kg, at 30-minute intervals) or saline (3 × 1 ml/kg, at 30-minute intervals).

### 2.6. Serum cocaine, benzoylecgonine (BE) and ecgonine methyl ester (EME) concentrations

Average serum cocaine concentration in rats treated with cocaine alone was 3.55 ± 0.48 µg/ml (mean ± SEM; n=8). In rats treated with haloperidol and cocaine, it was non-significantly higher, 6.34 ± 2.35 µg/ml (mean ± SEM; n=5). EME was detected in five of the eight rats treated with cocaine alone (0.96 ± 0.3 µg/ml) and in one of the five rats treated with haloperidol and cocaine. BE was not detected in rats treated with cocaine alone nor in rats treated with haloperidol and cocaine.

## 3. Discussion

At the behavioral level, cocaine evoked pronounced locomotor responses and stereotypic behaviors, which had been completely blocked after pretreatment with haloperidol. In the peripheral blood, haloperidol and cocaine had similar effects on immune cell numbers. Both drugs, whether administered separately or together, increased plasma corticosterone and evoked lymphopenia. The exception was that haloperidol prevented the cocaine-induced decrease in the numbers of NKT cells. Cocaine treatment altered the ratio of lymphocytes by increasing the percentage of T CD4^+^ cells while decreasing the percentage of B cells. Pretreatment with haloperidol did not change this effect. In the spleen, the infusion of cocaine after haloperidol significantly reduced the numbers of T CD4^+^ and B lymphocytes compared to rats treated with cocaine alone. This observation suggests a significant role for the D2-like DA receptors in retaining T CD4^+^ and B lymphocytes in the spleen after intravenous infusions of cocaine.

Cocaine activates the HPA axis [35], one of the major pathways through which cocaine modulates the immune system [19,20]. Haloperidol has bidirectional effects on the HPA axis, depending on the dose and route of administration: there is a tendency to increase the release of “stress hormones” at higher doses and routes, which provides a more rapid delivery of the drug to the brain [36,37]. HPA axis activation is commonly associated with lymphopenia and immunosuppression [38,39]. In our study, intravenous infusions of haloperidol and cocaine resulted in an increased concentration of plasma corticosterone, accompanied by pronounced lymphopenia. Similar effects were observed in rats receiving a high dose of exogenous corticosterone [40]. Surprisingly, in our study, haloperidol prevented the decrease in numbers of NKT cells caused by cocaine infusions. This result suggests that blocking D2-like DA receptors allows NKT cells to escape cocaine-induced lymphopenia. Peripheral DA was found to suppress the function of invariant NKT cells in the liver, however, through the pathway of the D1-like DA receptor-Protein Kinase A (PKA) [41].

One relevant result is that pronounced lymphopenia is coupled with an increased release of corticosterone as induced by haloperidol. The immunosuppressive effects of haloperidol have been reported, though they were missing important information regarding HPA axis activity [1,8,9]. Moreover, in 2007, the Food and Drug Administration (FDA) strengthened label warnings for intravenous haloperidol regarding its cardiac side effects. [42]. Our results support the emerging data that intravenous haloperidol in high doses is associated with the risk of side effects, including immunosuppression.

As observed in this study, cocaine-induced lymphopenia is associated with the increase in the proportion of T CD4^+^ cells. This effect was not mediated through DA receptors. In our previous experiments we found that the proportion of T CD4^+^ cells was increased on the 1^st^, 7^th^, and 14^th^ day of binge intravenous administration of cocaine despite the attenuation of HPA axis activation on the 7^th^ and 14^th^ day of treatment [34]. It is not clear which of the cocaine action mechanisms are responsible for this effect. This increase in the proportion of T CD4^+^ cells becomes more remarkable since there is evidence that alterations in the balance and function of T CD4^+^ cells may change the course of human immunodeficiency virus infections in cocaine users [43,44].

Reduced numbers of all tested lymphocyte subsets in peripheral blood may result from migration to peripheral lymphoid tissues or from cell death. Lymphocytes constantly migrate between peripheral blood and lymphatic tissues, and environmental cues can modulate this process [45,46]. In rats, in physiological conditions, mature lymphocytes migrate predominantly to the spleen or lymph nodes [47,48]. However, when environmental conditions are in flux (for example, when a rat is being exposed to social defeat stress), lymphocytes may change direction and travel to the bone marrow [49]. In our study, cocaine infusions induced lymphopenia in peripheral blood but at the same time showed a tendency to increase the numbers of B and T CD4^+^ lymphocytes in the spleen. This effect was reversed after pretreatment with haloperidol. When rats received haloperidol and subsequent cocaine infusions, we observed lymphopenia in both the peripheral blood and the spleen. Our results suggest that a DA signaling increase after cocaine treatment may be one of the mechanisms by which lymphocytes are retained in the spleen.

On the other hand, glucocorticoids that are increased by both cocaine and haloperidol (in addition to their effects on immune cells migration), also are known to induce lymphocyte apoptosis [50–52]. Cocaine has been shown to induce apoptosis of mice thymocytes *in vitro* [53] but the effects of both cocaine and haloperidol on apoptosis of blood or splenic lymphocytes *in vivo* are currently unknown [54].

Serum concentrations of IL-4 and IFN-γ both decreased after cocaine and haloperidol treatment; however, due to considerable inter-individual variability, we did not find the effects to be statistically significant. Results suggest that, for some individuals, haloperidol and cocaine may suppress IL-4 and IFN-γ signaling [18,55]. Moreover, the concentrations of IL-4 and IFN-γ that we detected in our study were comparable to that of control groups in other reports [56,57], suggesting that manipulations of chronically implanted catheters *per se* did not induce pronounced inflammation.

In conclusion, our results suggest that the binge pattern of administration of cocaine increases the proportion of T CD4^+^ lymphocytes in peripheral blood despite lymphopenia and elevated corticosterone. Blocking D2-like DA receptors allows NKT cells to escape the cocaine-induced lymphopenic effect. The DA signaling, which is elevated after cocaine administration, is involved in the retention of B and T CD4^+^ lymphocytes in the spleen. Finally, our results support the notion that intravenous haloperidol is associated with the risk of side effects, including immunosuppression.

## 4. Materials and methods

### 4.1. Animals

Thirty-two adult male Wistar rats (R. Grabowski Licensed Laboratory Animals Breeding, Gdansk, Poland) weighing 260-310 g were used. Upon arrival, animals were housed individually and handled by the experimenter daily for two weeks before experiments. Rats were kept in a temperature and humidity-controlled animal facility and maintained on a 12-h light/dark cycle (lights on from 06:00 to 18:00 h). They had free access to water and standard rodent food pellets except for behavioral sessions, including haloperidol and cocaine infusions. Animals were divided randomly into four groups of eight rats each. All procedures were approved by the Local Ethical Committee for the Care and Use of Laboratory Animals at the Medical University of Gdansk, Poland (No 34/2008).

### 4.2. Jugular vein cannulation and recovery after surgery

Rats were anesthetized with pentobarbital (60 mg/kg i.p., Vetbutal, Biowet Pulawy, Poland) and implanted with chronic Silastic catheters (i.d. 0.51 mm × o.d. 0.94 mm, Standard Silicone Tubing, HelixMark, Carpinteria, CA, USA) into the right jugular vein [34]. To prevent postoperative pain, rats received during the surgery subcutaneous injection of Carprofen 50 mg/ml (5% W/V) in a dose of about 12 mg/kg (Acticarp, Biowet Pulawy, Poland). Rats recovered after the implantation for at least 21 days before experimentation. During the first four days after the surgery, every 12 hours, each rat received 9.5 mg of ampicillin (Ampicillin 500 mg, Polfa Tarchomin S.A., Poland) dissolved in 0.3 ml of 0.9% sterile saline (Polpharma S.A., Poland) to prevent possible infections. The catheters were then flushed, every 12 hours, with a sterile 80 IU/ml solution of heparin in the volume of 0.3 ml (Heparinum natricum 5000 IU/ml inj., Warsaw Pharmaceutical Works Polfa, Poland) and then filled with 0.3 ml sterile 8000 IU/ml solution of streptase (Streptase 1 500 000 IU, ZLB Behring GmbH, Marburg, Germany). Beginning on the 7^th^ day after the surgery, catheters were flushed every 24 hours only with 0.4 ml of 0.9% sterile saline. All solutions used during the experiment were prepared in a laminar flow unit using sterilized plastic test tubes. All equipment used for infusions was sterile or disinfected in 70% ethanol for at least 12 hours before use.

### 4.3. Drugs and treatment

Haloperidol (Haloperidol 5 mg/ml, Warsaw Pharmaceutical Works Polfa, Poland) was diluted with sterile water for injections (Polpharma S.A., Poland) to obtain the concentration of 1 mg/ml. Haloperidol (1 mg/kg) or water for injections (1 ml/kg) were infused intravenously in freely moving rats 10 minutes before the first cocaine or saline infusion. The drugs were administered through a chronic jugular vein catheter connected with the PE tube which free end was outside the plexiglass chamber (Figure 1A and 1B). Immediately after pretreatment with haloperidol, the catheter was flushed with 0.3 ml of sterile water for injections. The dose of haloperidol was selected based on the potent inhibition of cocaine-induced locomotion in the preliminary dose-response test (Figure S1).

Cocaine hydrochloride (Sigma-Aldrich, St. Louis, MO, USA) was dissolved in 0.9% sterile saline (5 mg/ml). Sterile saline was used for control infusions. Each rat received three intravenous infusions of cocaine (3 × 5 mg/kg) or saline (3 × 1 ml/kg) at 30-minute intervals (Figure 1A). After the third infusion of cocaine, the catheter was flushed with 0.3 ml of sterile saline.

Haloperidol and cocaine were prepared immediately before use, and all solutions were administered in a volume of 1 ml/kg. Each haloperidol, cocaine, or control infusion lasted 30 seconds. All equipment used for infusions was sterile or disinfected in 70% ethanol for at least 12 hours before use.

### 4.4. Behavior

All infusions were made in freely moving rats placed in plexiglass chambers (43 × 43 × 20 cm) for measuring locomotor activity [34] (Figure 1B, Columbus Instruments, Opto-Varimex Minor, Columbus, OH, USA). For four consecutive days preceding haloperidol and cocaine administration, rats were placed in the chambers for 100-minute long sessions to habituate them to the new environment and to establish baseline locomotor activity assessed by the number of breaks in the horizontal infrared beams (data not shown). On the next day following the habituation period, rats were placed in the chambers and subjected to haloperidol and subsequent cocaine infusions. During the habituation sessions and on the experimental day, each rat was placed in the same chamber at the same time (between 8:00 and 14:00 h). Between experimental sessions, chambers were carefully cleaned with 70% ethanol.

### 4.5. Blood and spleen sampling

Blood samples were collected through a catheter 30 minutes after the third cocaine infusion. First, blood samples (0.5 ml) for determination of leukocyte numbers, cytofluorographic analysis, blood smears, and plasma were drawn using a sterilized polyethylene tube connected to a syringe containing 20 μl of 10% EDTA. Subsequently, additional blood samples (5 ml) for serum preparation were drawn into a syringe without anticoagulants. Immediately after the blood sampling, each rat was infused with a lethal dose of pentobarbital (160 mg in 1 ml, i.v., Morbital, Biowet Pulawy, Poland) administered through the catheter. Heartbeat and breath stopped within 2 - 5 s from the beginning of the infusion. Spleens were immediately harvested and placed in glass test tubes containing 5 ml of Hanks’ Balanced Salt Solution (HBSS, Sigma-Aldrich, St. Louis, MO, USA) with sodium bicarbonate (Merck, USA).

### 4.6. Preparation of spleen cells

Spleen suspended in 5 ml of HBSS was gently homogenized in a glass homogenizer. The obtained suspension was gently mixed and filtered through a nylon filter to separate splenocytes from the reticular connective tissue. Immediately after homogenization, the suspension of splenocytes was refrigerated (2 - 8°C) and analyzed within one hour.

### 4.7. Leukocytes and leukocyte subpopulations

Peripheral blood (10 μl) was diluted 20-fold with Türk solution and suspension of splenocytes in HBSS (10 μl) was diluted 60-fold and leukocytes were counted twice in a Neubauer hemocytometer. Leukocyte subpopulations were assessed by microscopic examination of peripheral blood and splenocyte suspension smears stained in a centrifuge (Aerospray Slide Stainer, 7120 Wescor, USA) by the May-Grünwald and Giemsa methods. Two smears of each sample were prepared. Percentages of leukocyte subpopulations were assessed on 200 or 400 cells per blood or spleen smear, respectively. Total numbers of lymphocytes and neutrophils were calculated by multiplying the total leukocyte number and percentage of lymphocytes or neutrophils.

### 4.8. Cytofluorographic analysis

Before samples preparation, suspension of splenocytes (30 µl) was diluted with HBSS (300 µl) to obtain an optimal frequency of cells detection during cytofluorographic analysis. Three-color flow cytometry of whole blood samples and diluted suspension of splenocytes was performed using the following antibodies: IOTest CD3-FITC/CD45RA-PC7/CD161a-APC and CD3-FITC/CD4-PC7/CD8-APC (Beckman Coulter, Immunotech, Marseille, France) for determination of T, B, NK, NKT, and T CD4^+^ and T CD8^+^ lymphocytes, respectively. In the present study, rat lymphocytes with co-expression of CD3 and CD161a molecules were classified as the NKT cells [58]. Cells that express on their surface both the CD3 molecules and a diverse set of NK cell markers (e.g., CD161, CD16 or CD56) are described as NKT cells [59,60]. According to the manufacturer’s instructions, each antibody (25 μl) was added to the separate whole blood samples (25 μl) or a diluted suspension of splenocytes (25 μl), mixed and incubated for 20 minutes in the darkness. After the first incubation, erythrocytes were lysed (1 ml of Versalyse, Beckman Coulter) and the lymphocytes fixed (25 μl of Fixative Solution, Beckman Coulter). The samples were mixed and incubated for 10 minutes in the darkness. Flow cytometry was performed in a model FC500 cytometer (Beckman Coulter, Inc., Brea, CA, USA). During the assay, the lymphocytes were gated and percentages of tested cell phenotypes were assessed.

### 4.9. Plasma corticosterone measurement

Blood samples (0.4 ml, containing 16 μl of 10% EDTA) were centrifuged. Plasma samples were collected and immediately stored at -70°C until testing. Quantitative determination of corticosterone concentration in plasma was performed in duplicates in a single assay using a commercial radioimmunoassay kit (DRG Diagnostics GmbH, Marburg, Germany) and Wizard 1470 gamma counter (Pharmacia LKB, Turku, Finland). The assay was performed according to the manufacturer’s instructions.

### 4.10. IL-4 and IFN-γ measurements

Blood samples (5 ml) were left for clotting at room temperature for 20 minutes and centrifuged. The serum samples for measuring concentrations of IL-4 and IFN-γ were collected and immediately stored at -70°C until testing. Quantitative determinations of IL-4 and IFN-γ concentrations in rat serum were performed in duplicates in a single assay for IL-4 or IFN-γ, using a specific commercial cytokine ELISA kits (GEN-PROBE Diaclone, Besancon, France). The assay was performed according to the manufacturer’s instructions. IL-4 and IFN-γ concentrations were determined using DTX 880 Multimode Detector (Beckman Coulter, Inc., Brea, CA, USA) set to 450 nm.

### 4.11. Cocaine, benzoylecgonine (BE) and ecgonine methyl ester (EME) determinations

Cocaine, BE and EME were detected and determined by gas chromatography coupled with a mass spectrometry detector (GC-MS). Blood samples (5 ml) were left for clotting at room temperature for 20 minutes and centrifuged. The serum samples for measuring concentrations of cocaine, BE and EME were collected and immediately stored at -70°C. Upon testing, the mass of serum samples was determined and the pH of the samples was adjusted to pH 9 using the ammonium buffer (NH4Cl/NH3, pH 9.5). The samples were placed on the columns packed with diatomaceous earth (Extrelut NT 1, Merck, Darmstadt, Germany) and left for 20 minutes at room temperature. Elution of analytes was performed with 5 ml of dichloromethane and isopropanol mixture (85:15). Eluate was evaporated to dryness in the mild stream of nitrogen at the temperature of 40°C. The dried residue was dissolved in 50 μl of acetonitrile and 2 μl of the prepared sample extract was injected into the GC-MS set for analysis. Gas chromatograph TRACE GC coupled with mass spectrometry detector TRACE DSQ (ThermoFinnigan, Milan, Italy) was used for the analysis under the following conditions: capillary column, Zebron ZB-5 Guardian (30 m × 0.25 mm i.d. × 0.25 µm film thickness, Phenomenex, Torrance, CA, USA); split-splitless capillary inlet system was operated in split mode at 280°C; chromatograph oven temperature range 50-285°C; carrier gas, helium. Mass spectrometry detector was operated in the SIM mode using the following molecular ions for quantification: cocaine, *m/z* 82, 182, 77; benzoylecgonine, *m/z* 82, 124, 168; ecgonine methyl ester, *m/z* 82, 96, 168. Detection levels for cocaine, EME, and BE were 1.5 µg/ml, 0.4 µg/ml, and 13 µg/ml, respectively.

### 4.12. Statistical analysis

Two-way repeated-measures analysis of variance (ANOVA), with cocaine and haloperidol as factors, was used to analyze data. Post-hoc comparisons were performed using Student’s t-test with Bonferroni correction. One-way ANOVA was used to compare the cocaine and cocaine metabolites concentrations between the groups.

## Author Contributions

Conceptualization, MMJ and AHS; Methodology, MMJ, Formal analysis, MMJ; Investigation, MMJ, BMJ, WG, MW; Data curation, MMJ; Writing - original draft preparation, MMJ, BMJ, PKG and AHS; Supervision, AHS; Project administration, MMJ; Funding acquisition, MMJ, BMJ and AHS. All authors have read and agreed to the published version of the manuscript.

## Institutional Review Board Statement

All surgical procedures and experimental protocols were performed with the Guide for Using Animal Subjects, and approved by the Local Ethical Committee for the Care and Use of Laboratory Animals at the Medical University of Gdansk, Poland (No 34/2008).

## Data Availability Statement

The data presented in this study will be provided upon request.

## Acknowledgments

We thank the Polish Ministry of Science and Higher Education for funding Research Grants N N303 417137 and N N303 394036 to M.M. Jankowski and B. Ignatowska-Jankowska.

---

**Figure S1.**
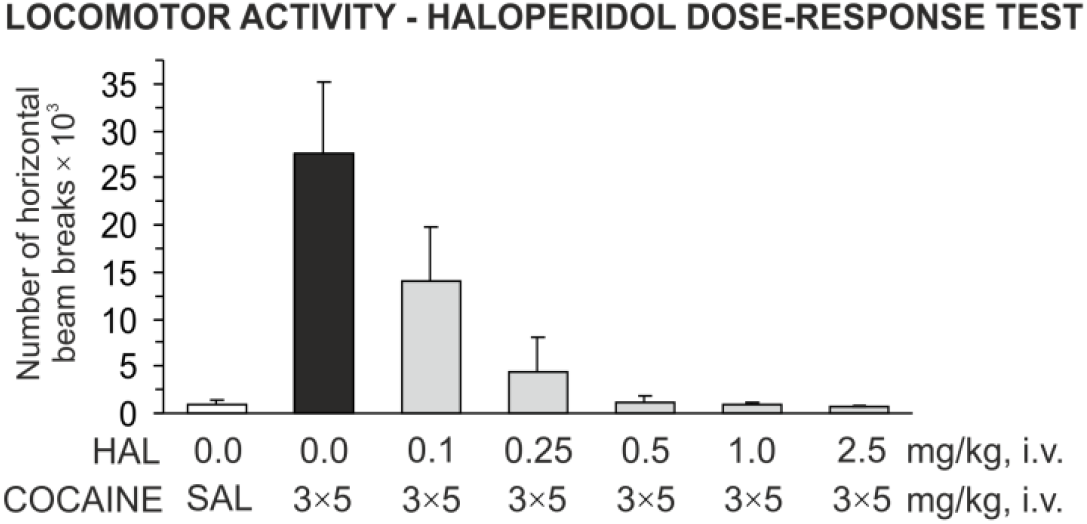
Inhibition of cocaine-induced locomotor activity by haloperidol. Haloperidol (HAL) in doses of 0.1 mg/kg, 0.25 mg/kg; 0.5 mg/kg; 1.0 mg/kg, 2.5 mg/kg was administered i.v. 10 minutes before the first infusion of either saline (SAL; 3 × 1 ml/kg) or cocaine (3 × 5 mg/kg) infusion. Figure shows mean ± SD.

## References

1. Lao, K.S.J.; Wong, A.Y.S.; Wong, I.C.K.; Besag, F.M.C.; Chang, W.C.; Lee, E.H.M.; Chen, E.Y.H.; Blais, J.E.; Chan, E.W. Mortality Risk Associated with Haloperidol Use Compared with Other Antipsychotics: An 11-Year Population-Based Propensity-Score-Matched Cohort Study. CNS Drugs 2020, 34, 197–206, doi:10.1007/s40263-019-00693-5.

2. Gómez-Revuelta, M.; Pelayo-Terán, J.M.; Juncal-Ruiz, M.; Vázquez-Bourgon, J.; Suárez-Pinilla, P.; Romero-Jiménez, R.; Setién Suero, E.; Ayesa-Arriola, R.; Crespo-Facorro, B. Antipsychotic Treatment Effectiveness in First Episode of Psychosis: PAFIP 3-Year Follow-Up Randomized Clinical Trials Comparing Haloperidol, Olanzapine, Risperidone, Aripiprazole, Quetiapine, and Ziprasidone. International Journal of Neuropsychopharmacology 2020, 23, 217–229, doi:10.1093/ijnp/pyaa004.

3. Martel, J.C.; Gatti McArthur, S. Dopamine Receptor Subtypes, Physiology and Pharmacology: New Ligands and Concepts in Schizophrenia. Front. Pharmacol. 2020, 11, 1003, doi:10.3389/fphar.2020.01003.

4. Marwari, S.; Dawe, G.S. Effects of Haloperidol on Cognitive Function and Behavioural Flexibility in the IntelliCage Social Home Cage Environment. Behavioural Brain Research 2019, 371, 111976, doi:10.1016/j.bbr.2019.111976.

5. Waku, I.; Magalhães, M.S.; Alves, C.O.; de Oliveira, A.R. Haloperidol-induced Catalepsy as an Animal Model for Parkinsonism: A Systematic Review of Experimental Studies. Eur J Neurosci 2021, 53, 3743–3767, doi:10.1111/ejn.15222.

6. Green, M.F.; Marder, S.R.; Glynn, S.M.; McGurk, S.R.; Wirshing, W.C.; Wirshing, D.A.; Liberman, R.P.; Mintz, J. The Neurocognitive Effects of Low-Dose Haloperidol: A Two-Year Comparison with Risperidone. Biological Psychiatry 2002, 51, 972–978, doi:10.1016/S0006-3223(02)01370-7.

7. Schmack, K.; Bosc, M.; Ott, T.; Sturgill, J.F.; Kepecs, A. Striatal Dopamine Mediates Hallucination-like Perception in Mice. Science 2021, 372, eabf4740, doi:10.1126/science.abf4740.

8. Leykin, I.; Mayer, R.; Shinitzky, M. Short and Long Term Immunosuppressive Effects of Clozapine and Haloperidol. Immunopharmacology 1997, 37, 75–86, doi:10.1016/S0162-3109(97)00037-4.

9. Song, C.; Lin, A.; Kenis, G.; Bosmans, E.; Maes, M. Immunosuppressive Effects of Clozapine and Haloperidol: Enhanced Production of the Interleukin-1 Receptor Antagonist. Schizophrenia Research 2000, 42, 157–164, doi:10.1016/S0920-9964(99)00116-4.

10. Al-Amin, M.; Uddin, M.M.N.; Reza, H.M. Effects of Antipsychotics on the Inflammatory Response System of Patients with Schizophrenia in Peripheral Blood Mononuclear Cell Cultures. Clin Psychopharmacol Neurosci 2013, 11, 144–151, doi:10.9758/cpn.2013.11.3.144.

11. Yamamoto, S.; Ohta, N.; Matsumoto, A.; Horiguchi, Y.; Koide, M.; Fujino, Y. Haloperidol Suppresses NF-KappaB to Inhibit Lipopolysaccharide-Induced Pro-Inflammatory Response in RAW 264 Cells. Med Sci Monit 2016, 22, 367–372, doi:10.12659/MSM.895739.

12. Cotel, M.-C.; Lenartowicz, E.M.; Natesan, S.; Modo, M.M.; Cooper, J.D.; Williams, S.C.R.; Kapur, S.; Vernon, A.C. Microglial Activation in the Rat Brain Following Chronic Antipsychotic Treatment at Clinically Relevant Doses. European Neuropsychopharmacology 2015, 25, 2098–2107, doi:10.1016/j.euroneuro.2015.08.004.

13. Bloomfield, P.S.; Bonsall, D.; Wells, L.; Dormann, D.; Howes, O.; Paola, V.D. The Effects of Haloperidol on Microglial Morphology and Translocator Protein Levels: An in Vivo Study in Rats Using an Automated Cell Evaluation Pipeline. J Psychopharmacol 2018, 32, 1264–1272, doi:10.1177/0269881118788830.

14. Racki, V.; Marcelic, M.; Stimac, I.; Petric, D.; Kucic, N. Effects of Haloperidol, Risperidone, and Aripiprazole on the Immunometabolic Properties of BV-2 Microglial Cells. IJMS 2021, 22, 4399, doi:10.3390/ijms22094399.

15. Pellegrino, T.; Bayer, B.M. In Vivo Effects of Cocaine on Immune Cell Function. Journal of Neuroimmunology 1998, 83, 139–147, doi:10.1016/S0165-5728(97)00230-0.

16. Pellegrino, T.C.; Dunn, K.L.; Bayer, B.M. Mechanisms of Cocaine-Induced Decreases in Immune Cell Function. International Immunopharmacology 2001, 1, 665–675, doi:10.1016/S1567-5769(00)00051-5.

17. Irwin, M.R.; Olmos, L.; Wang, M.; Valladares, E.M.; Motivala, S.J.; Fong, T.; Newton, T.; Butch, A.; Olmstead, R.; Cole, S.W. Cocaine Dependence and Acute Cocaine Induce Decreases of Monocyte Proinflammatory Cytokine Expression across the Diurnal Period: Autonomic Mechanisms. J Pharmacol Exp Ther 2007, 320, 507–515, doi:10.1124/jpet.106.112797.

18. Zaparte, A.; Schuch, J.B.; Viola, T.W.; Baptista, T.A.S.; Beidacki, A.S.; do Prado, C.H.; Sanvicente-Vieira, B.; Bauer, M.E.; Grassi-Oliveira, R. Cocaine Use Disorder Is Associated With Changes in Th1/Th2/Th17 Cytokines and Lymphocytes Subsets. Front. Immunol. 2019, 10, 2435, doi:10.3389/fimmu.2019.02435.

19. Stanulis, E.D.; Matulka, R.A.; Jordan, S.D.; Rosecrans, J.A.; Holsapple, M.P. Role of Corticosterone in the Enhancement of the Antibody Response after Acute Cocaine Administration. J Pharmacol Exp Ther 1997, 280, 284–291.

20. Stanulis, E.D.; Jordan, S.D.; Rosecrans, J.A.; Holsapple, M.P. Disruption of Th1/Th2 Cytokine Balance by Cocaine Is Mediated by Corticosterone. Immunopharmacology 1997, 37, 25–33, doi:10.1016/S0162-3109(96)00167-1.

21. Caroleo, M.C.; Arbirio, M.; Di Francesco, P.; Pulvirenti, L.; Garaci, E.; Nistico, G. Cocaine Induced T Cell Proliferation in the Rat: Role of Amygdala Dopamine D1 Receptors. Neuroscience Letters 1998, 256, 61–64, doi:10.1016/S0304-3940(98)00758-7.

22. Levite, M. Dopamine and T Cells: Dopamine Receptors and Potent Effects on T Cells, Dopamine Production in T Cells, and Abnormalities in the Dopaminergic System in T Cells in Autoimmune, Neurological and Psychiatric Diseases. Acta Physiol 2016, 216, 42–89, doi:10.1111/apha.12476.

23. Matt, S.M.; Gaskill, P.J. Where Is Dopamine and How Do Immune Cells See It?: Dopamine-Mediated Immune Cell Function in Health and Disease. J Neuroimmune Pharmacol 2020, 15, 114–164, doi:10.1007/s11481-019-09851-4.

24. Cosentino, M.; Ferrari, M.; Kustrimovic, N.; Rasini, E.; Marino, F. Influence of Dopamine Receptor Gene Polymorphisms on Circulating T Lymphocytes: A Pilot Study in Healthy Subjects. Human Immunology 2015, 76, 747–752, doi:10.1016/j.humimm.2015.09.032.

25. Cui, Y.; Prabhu, V.; Nguyen, T.; Yadav, B.; Chung, Y.-C. The MRNA Expression Status of Dopamine Receptor D2, Dopamine Receptor D3 and DARPP-32 in T Lymphocytes of Patients with Early Psychosis. IJMS 2015, 16, 26677–26686, doi:10.3390/ijms161125983.

26. McKenna, F.; McLaughlin, P.J.; Lewis, B.J.; Sibbring, G.C.; Cummerson, J.A.; Bowen-Jones, D.; Moots, R.J. Dopamine Receptor Expression on Human T- and B-Lymphocytes, Monocytes, Neutrophils, Eosinophils and NK Cells: A Flow Cytometric Study. Journal of Neuroimmunology 2002, 132, 34–40, doi:10.1016/S0165-5728(02)00280-1.

27. Tomassoni, D.; Bronzetti, E.; Cantalamessa, F.; Mignini, F.; Ricci, A.; Sabbatini, M.; Tayebati, S.K.; Zaccheo, D. Postnatal Development of Dopamine Receptor Expression in Rat Peripheral Blood Lymphocytes. Mechanisms of Ageing and Development 2002, 123, 491–498, doi:10.1016/S0047-6374(01)00355-4.

28. Qiu, Y.-H.; Peng, Y.-P.; Jiang, J.-M.; Wang, J.-J. Expression of Tyrosine Hydroxylase in Lymphocytes and Effect of Endogenous Catecholamines on Lymphocyte Function. Neuroimmunomodulation 2004, 11, 75–83, doi:10.1159/000075316.

29. Marazziti, D.; Consoli, G.; Masala, I.; Catena Dell’Osso, M.; Baroni, S. Latest Advancements on Serotonin and Dopamine Transporters in Lymphocytes. MRMC 2010, 10, 32–40, doi:10.2174/138955710791112587.

30. Mignini, F.; Sabbatini, M.; Capacchietti, M.; Amantini, C.; Bianchi, E.; Artico, M.; Tammaro, A. T-Cell Subpopulations Express a Different Pattern of Dopaminergic Markers in Intra- and Extra-Thymic Compartments. J Biol Regul Homeost Agents 2013, 27, 463–475.

31. Benesics, A.; Sershen, H.; Baranyi, M.; Hashim, A.; Lajtha, A.; Sylvester Vizi, E. Dopamine, as Well as Norepinephrine, Is a Link between Noradrenergic Nerve Terminals and Splenocytes. Brain Research 1997, 761, 236–243, doi:10.1016/S0006-8993(97)00313-2.

32. Mignini, F.; Traini, E.; Tomassoni, D.; Amenta, F. Dopamine Plasma Membrane Transporter (DAT) in Rat Thymus and Spleen: An Immunochemical and Immunohistochemical Study. Autonom & Auta Pharm 2006, 26, 183–189, doi:10.1111/j.1474-8673.2006.00370.x.

33. Mignini, F.; Tomassoni, D.; Traini, E.; Amenta, F. Dopamine, Vesicular Transporters and Dopamine Receptor Expression and Localization in Rat Thymus and Spleen. Journal of Neuroimmunology 2009, 206, 5–13, doi:10.1016/j.jneuroim.2008.09.018.

34. Jankowski, M.M.; Ignatowska-Jankowska, B.; Glac, W.; Swiergiel, A.H. Cocaine Administration Increases CD4/CD8 Lymphocyte Ratio in Peripheral Blood despite Lymphopenia and Elevated Corticosterone. International Immunopharmacology 2010, 10, 1229–1234, doi:10.1016/j.intimp.2010.07.003.

35. Manetti, L.; Cavagnini, F.; Martino, E.; Ambrogio, A. Effects of Cocaine on the Hypothalamic–Pituitary–Adrenal Axis. J Endocrinol Invest 2014, 37, 701–708, doi:10.1007/s40618-014-0091-8.

36. Lourenço, G.A.; Dorce, V.A.C.; Palermo-Neto, J. Haloperidol Treatments Increased Macrophage Activity in Male and Female Rats: Influence of Corticosterone and Prolactin Serum Levels. European Neuropsychopharmacology 2005, 15, 271–277, doi:10.1016/j.euroneuro.2004.11.007.

37. Handley, R.; Mondelli, V.; Zelaya, F.; Marques, T.; Taylor, H.; Reinders, A.A.T.S.; Chaddock, C.; McQueen, G.; Hubbard, K.; Papadopoulos, A.; et al. Effects of Antipsychotics on Cortisol, Interleukin-6 and Hippocampal Perfusion in Healthy Volunteers. Schizophrenia Research 2016, 174, 99–105, doi:10.1016/j.schres.2016.03.039.

38. Chen, A.; Sun, X.; Wang, W.; Liu, J.; Zeng, X.; Qiu, J.; Liu, X.; Wang, Y. Activation of the Hypothalamic-Pituitary-Adrenal (HPA) Axis Contributes to the Immunosuppression of Mice Infected with Angiostrongylus Cantonensis. J Neuroinflammation 2016, 13, 266, doi:10.1186/s12974-016-0743-z.

39. Zierath, D.; Tanzi, P.; Shibata, D.; Becker, K.J. Cortisol Is More Important than Metanephrines in Driving Changes in Leukocyte Counts after Stroke. Journal of Stroke and Cerebrovascular Diseases 2018, 27, 555–562, doi:10.1016/j.jstrokecerebrovasdis.2017.09.048.

40. Flaherty, D.K.; McGarity, K.L.; Winzenburger, P.; Panyik, M. The Effect of Continuous Corticosterone Administration On Lymphocyte Subpopulations in the Peripheral Blood of the Fischer 344 Rat as Determined by Two Color Flow Cytometric Analyses. Immunopharmacology and Immunotoxicology 1993, 15, 583–604, doi:10.3109/08923979309019732.

41. Xue, R.; Zhang, H.; Pan, J.; Du, Z.; Zhou, W.; Zhang, Z.; Tian, Z.; Zhou, R.; Bai, L. Peripheral Dopamine Controlled by Gut Microbes Inhibits Invariant Natural Killer T Cell-Mediated Hepatitis. Front. Immunol. 2018, 9, 2398, doi:10.3389/fimmu.2018.02398.

42. Meyer-Massetti, C.; Cheng, C.M.; Sharpe, B.A.; Meier, C.R.; Guglielmo, B.J. The FDA Extended Warning for Intravenous Haloperidol and Torsades de Pointes: How Should Institutions Respond? J. Hosp. Med.2010, 5, E8–E16, doi:10.1002/jhm.691.

43. Mantri, C.K.; Pandhare Dash, J.; Mantri, J.V.; Dash, C.C.V. Cocaine Enhances HIV-1 Replication in CD4+ T Cells by Down-Regulating MiR-125b. PLoS ONE 2012, 7, e51387, doi:10.1371/journal.pone.0051387.

44. Pandhare, J.; Addai, A.B.; Mantri, C.K.; Hager, C.; Smith, R.M.; Barnett, L.; Villalta, F.; Kalams, S.A.; Dash, C. Cocaine Enhances HIV-1–Induced CD4+ T-Cell Apoptosis. The American Journal of Pathology 2014, 184, 927–936, doi:10.1016/j.ajpath.2013.12.004.

45. Kindt, T.J.; Goldsby, R.A.; Osborne, B.A.; Kuby, J. Kuby Immunology; 6. ed.; Freeman: New York, 2007; ISBN 978-0-7167-8590-3.

46. Vuononvirta, J.; Marelli-Berg, F.M.; Poobalasingam, T. Metabolic Regulation of T Lymphocyte Motility and Migration. Molecular Aspects of Medicine 2021, 77, 100888, doi:10.1016/j.mam.2020.100888.

47. Johnsson, C.; Festin, R.; Tufveson, G.; Tötterman, T.H. Ex Vivo PKH26-Labelling of Lymphocytes for Studies of Cell Migration in Vivo. Scand J Immunol 1997, 45, 511–514, doi:10.1046/j.1365-3083.1997.d01-430.x.

48. Fulgenzi, A.; Ferrero, E.; Gasparini, M.; Casati, R.; Riccardo Colombo, F.; Gerundini, P.; Elena Ferrero, M. Technetium-99m Scintigraphy to Visualize T-Cell Homing in Vivo: A Preclinical Study. Nuclear Medicine and Biology 2003, 30, 633–642, doi:10.1016/S0969-8051(03)00051-9.

49. Stefanski, V.; Peschel, A.; Reber, S. Social Stress Affects Migration of Blood T Cells into Lymphoid Organs. Journal of Neuroimmunology 2003, 138, 17–24, doi:10.1016/S0165-5728(03)00076-6.

50. Yao, Z.; DuBois, D.C.; Almon, R.R.; Jusko, W.J. Pharmacokinetic/Pharmacodynamic Modeling of Corticosterone Suppression and Lymphocytopenia by Methylprednisolone in Rats. Journal of Pharmaceutical Sciences 2008, 97, 2820–2832, doi:10.1002/jps.21167.

51. Cain, D.W.; Cidlowski, J.A. Immune Regulation by Glucocorticoids. Nat Rev Immunol 2017, 17, 233–247, doi:10.1038/nri.2017.1.

52. Shimba, A.; Ikuta, K. Control of Immunity by Glucocorticoids in Health and Disease. Semin Immunopathol 2020, 42, 669–680, doi:10.1007/s00281-020-00827-8.

53. Wu, Y.-B.; Shen, M.-L.; Gu, G.-G.; Anderson, K.M.; Ou, D.W. The Effects of Cocaine Injections on Mouse Thymocyte Population. Experimental Biology and Medicine 1997, 214, 173–179, doi:10.3181/00379727-214-44085.

54. Northcutt, A.L.; Hutchinson, M.R.; Wang, X.; Baratta, M.V.; Hiranita, T.; Cochran, T.A.; Pomrenze, M.B.; Galer, E.L.; Kopajtic, T.A.; Li, C.M.; et al. DAT Isn’t All That: Cocaine Reward and Reinforcement Require Toll-like Receptor 4 Signaling. Mol Psychiatry 2015, 20, 1525–1537, doi:10.1038/mp.2014.177.

55. Stamatovich, S.N.; Lopez-Gamundi, P.; Suchting, R.; Colpo, G.D.; Walss-Bass, C.; Lane, S.D.; Schmitz, J.M.; Wardle, M.C. Plasma Pro- and Anti-Inflammatory Cytokines May Relate to Cocaine Use, Cognitive Functioning, and Depressive Symptoms in Cocaine Use Disorder. The American Journal of Drug and Alcohol Abuse 2021, 47, 52–64, doi:10.1080/00952990.2020.1828439.

56. Zheng, K.-C.; Nong, D.-X.; Morioka, T.; Todoriki, H.; Ariizumi, M. Elevated Interleukin-4 and Interleukin-6 in Rats Sensitized with Toluene Diisocyanate. INDUSTRIAL HEALTH 2001, 39, 334–339, doi:10.2486/indhealth.39.334.

57. Pertsov, S.S.; Koplik, E.V.; Stepanyuk, V.L.; Simbirtsev, A.S. Blood Cytokines in Rats with Various Behavioral Characteristics during Emotional Stress and Treatment with Interleukin-1β. Bull Exp Biol Med 2009, 148, 196–199, doi:10.1007/s10517-009-0668-y.

58. Ignatowska-Jankowska, B.; Jankowski, M.; Glac, W.; Swiergel, A.H. Cannabidiol-Induced Lymphopenia Does Not Involve NKT and NK Cells. J Physiol Pharmacol 2009, 60 Suppl 3, 99–103.

59. Mittag, A.; Lenz, D.; Gerstner, A.O.H.; Sack, U.; Steinbrecher, M.; Koksch, M.; Raffael, A.; Bocsi, J.; Tárnok, A. Polychromatic (Eight-Color) Slide-Based Cytometry for the Phenotyping of Leukocyte, NK, and NKT Subsets. Cytometry 2005, 65A, 103–115, doi:10.1002/cyto.a.20140.

60. Reis, E.A.G.; Athanazio, D.A.; Lima, I.; e Silva, N.O.; Andrade, J.C.S.; Jesus, R.N.; Barbosa, L.M.; Reis, M.G.; Santiago, M.B. NK and NKT Cell Dynamics after Rituximab Therapy for Systemic Lupus Erythematosus and Rheumatoid Arthritis. Rheumatol Int 2009, 29, 469–475, doi:10.1007/s00296-008-0719-0.

